# Conserved meiotic mechanisms in the cnidarian *Clytia hemisphaerica* revealed by Spo11 knockout

**DOI:** 10.1101/2022.01.05.475076

**Authors:** Catriona Munro, Hugo Cadis, Sophie Pagnotta, Evelyn Houliston, Jean-René Huynh

## Abstract

During meiosis, DNA recombination allows the shuffling of genetic information between the maternal and paternal chromosomes. Recombination is initiated by double strand breaks (DSBs) catalyzed by the conserved enzyme Spo11. How this crucial event is connected to other meiotic processes is surprisingly variable depending on the species. Here, we knocked down *Spo11* by CRISPR in the jellyfish *Clytia hemisphaerica*, belonging to Cnidaria, the sister group to Bilateria (where classical animal models are found). *Spo11* mutants fail to assemble synaptonemal complexes and chiasmata, and in consequence homologous chromosome pairs disperse during oocyte growth, creating aneuploid but fertilizable eggs that develop into viable larvae. *Clytia* thus shares an ancient eukaryotic dependence of synapsis and chromosome segregation on Spo11-generated DSBs, and provides new evolutionary perspectives on meiosis regulation.

## Introduction

In meiosis, diploid germ cells undergo two rounds of nuclear division to produce haploid gametes. A crucial feature of meiosis is the pairing of maternal and paternal chromosome homologs during an extended prophase, before segregating from each other on the first meiotic spindle (1, 2). Their initial loose coupling is stabilized by the polymerisation of a proteinaceous structure called the synaptonemal complex (SC), which holds together homologous axes and promotes genetic recombination (3). Exchanges of chromatids, or crossovers, allow for the formation of physical links (chiasmata) which maintain homologs associated in pairs following depolymerization of the SC. Individual meiotic processes and molecular machinery at each step are generally well conserved among eukaryotes and likely descend from a common eukaryotic ancestor (4). Surprisingly, however, functional studies in classical model organisms have revealed a surprising diversity in the chronology and interdependency of these early meiotic steps (1, 2). This paradox has been highlighted by studies of Spo11, a meiosis-specific enzyme highly conserved across eukaryotes that is required for the formation of DNA double-stranded breaks (DSBs) and the initiation of meiotic recombination (5–14). In many of the species studied, Spo11-generated DSBs are required for pairing and synapsis of the replicated homologous chromosomes, with crossover sites (chiasmata) required subsequently for maintaining the pairs until their correct segregation during meiotic divisions (5, 8–11, 14). There are, however, several exceptions, notably in the classical animal model species *Caenorhabditis elegans* and female *Drosophila melanogaster* (6, 7). All animal models in which Spo11 function has been examined so far belong to the clade Bilateria (Figure 1A). Cnidaria, which includes corals, anemones, and jellyfish and is the sister clade to Bilateria, offers a valuable evolutionary perspective on meiotic processes. The jellyfish *Clytia hemisphaerica* is now established as a cnidarian model species well suited for experimental manipulation, with the simple organization and transparency of the gonads making them attractive for analyzing gametogenesis regulation in vivo (15, 16). Furthermore, particularities of its life cycle (Figure 1B) conveniently enable phenotypes of gene mutations targeted by CRISPR-Cas9 to be studied in clonally-produced F0 generation jellyfish, which bud from stable, vegetatively-propagating polyp colonies (17).

**Figure 1:**
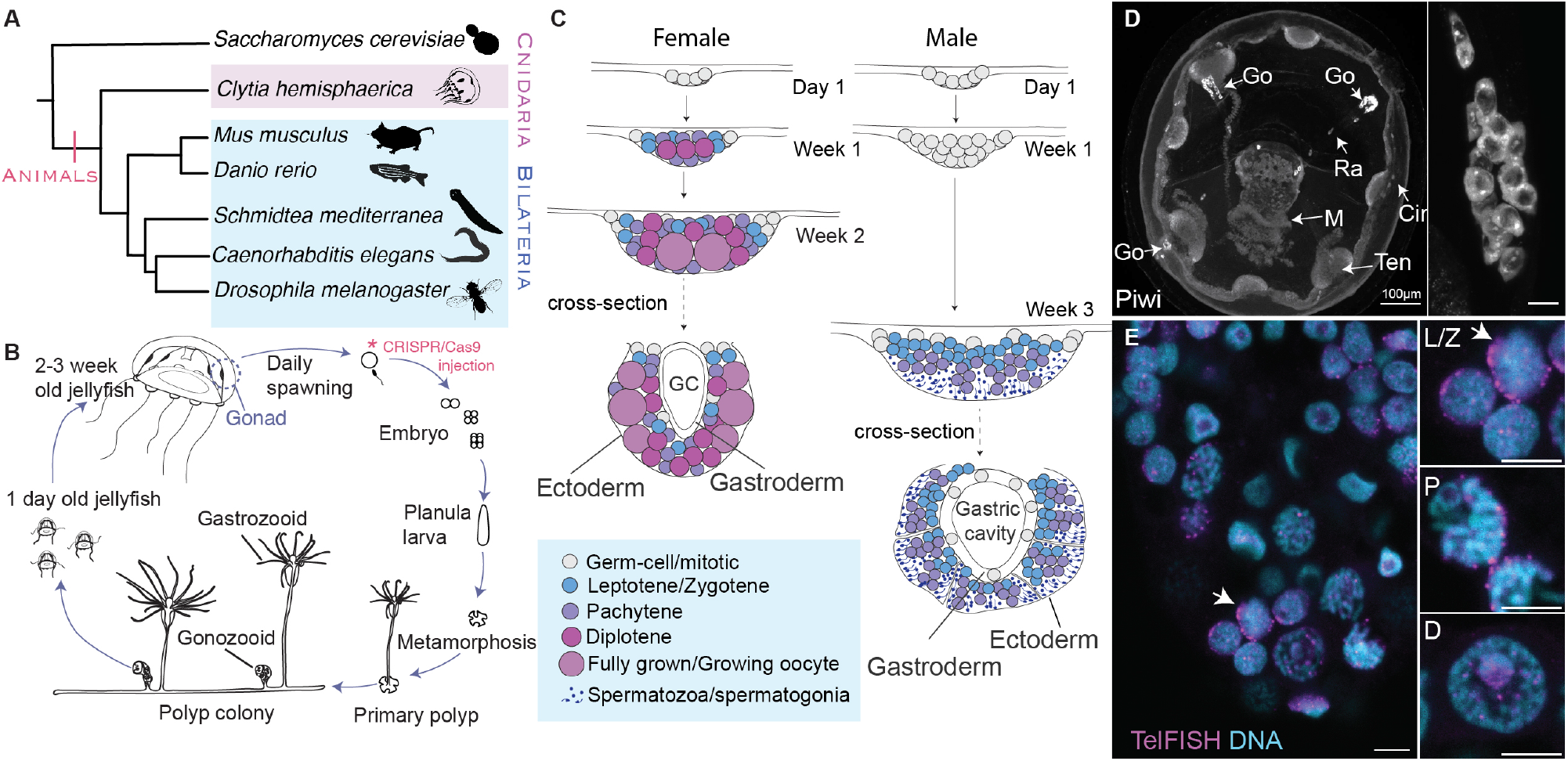
Phylogenetic placement, life-cycle, and gonad organization of *Clytia hemisphaerica*. A. Evolutionary relationships among major animal meiosis models. Silhouettes from PhyloPic.org, *S. mediterranea*: credit to Noah Schlottman, *C. hemisphaerica*: credit to Joseph Ryan, photo by Patrick Steinmetz, under a share-alike license https://creativecommons.org/licenses/by-sa/3.0/. B. Life-cycle of *Clytia hemisphaerica*. The polyp colony propagates asexually for many years in the lab. Specialized polyps called gonozooids release jellyfish continuously. Sexually mature jellyfish release eggs or sperm daily in response to dark-light transitions. The embryo develops into a “planula” larva which settles and metamorphoses to form a primary polyp. To achieve gene knockouts, CRISPR/Cas9 is injected in the mature egg immediately prior to fertilization. C. Development and organization of the male and female gonad from the first day of jellyfish release to sexual maturity. GC = gastric cavity. D. Maximum intensity projection of a confocal image stack of a three day old jellyfish stained with anti-Piwi. Go = gonad, Ra = radial canal, M = manubrium/mouth, Ten = tentacle bulb, Cir = circular canal. Scale is 100 µm. Similar projection from a 3 day old gonad is shown on the right, scale is 10 µm. E. Confocal section showing oocytes at mixed meiotic stages within the gonad of a one week female jellyfish stained with telomere FISH (magenta) and Hoechst (cyan). Inset shows close-up of representative nuclei at different prophase I stages, L/Z = leptotene/zygotene, P = pachytene, D = diplotene. Arrow indicates telomere bouquet. Scale is 5 µm.

### Mapping early meiosis in *Clytia* jellyfish

To establish a framework for studies of early meiotic mechanisms in *Clytia*, we first documented the formation of the gonads in male and female jellyfish. Following their release from the polyp colony baby jellyfish show four small patches of germ-line precursor cells enriched in Piwi protein (18) midway along each radial canal at the sites of the future gonads (Figure 1C, 1D, Supplementary Figure S1A,B). The gonads develop and expand as the jellyfish grows to sexual maturity, from about 1mm to 1cm in diameter over 2-3 weeks, remaining clearly visible within the transparent animal. They maintain a simple overall organization as germline cells proliferate and differentiate between two somatic tissue layers: the gastrodermis and the epidermis (Figure 1C). In females, germ cells entering meiosis are detected by 4-5 days, when oocytes of all stages of prophase I can be identified (Figure 1C, 1E). Hoechst staining reveals classic figures of condensed chromatin during leptotene-zygotene, and structured parallel tracks indicative of pairing and synapsis in pachytene (Figure 1E). Diplotene nuclei show a distinctive Hoechst-negative “hole” with one or more diffuse Hoecht-positive patches appearing as the oocyte grows (Figure 1E). Telomere FISH shows characteristic “bouquet” configurations in leptotene/zygotene oocytes (Figure 1E). Diplotene arrested oocytes undergo extensive growth to reach 180- 200µm in diameter. Fully grown oocytes become competent to undergo meiotic maturation and spawning, a process triggered daily in mature ovaries by a peptide hormone (MIH) secreted in response to light (19–21). Following each spawning, “stage I” growing oocytes of about 50µm in diameter grow to full size in around 13-15 hours (16, 21). Towards the end of this growth phase the chromosomes condense in preparation for diakinesis (16). *Clytia* male jellyfish undergo a longer period of proliferation of germ-line precursors than females (Figure 1C, Supplementary Figure S1C,D). After 2 weeks, prophase I spermatogonia are distributed in layers, with leptotene/zygotene cells closest to the gastrodermis, then pachytene, spermatids, and finally spermatozoa closest to the ectoderm (Figure 1C, Supplementary Figure S1c,d). Daily sperm release occurs approximately 1.5 h after light exposure. Within each subsequent 24 hour period, cohorts of spermatogonia enter and progress through pachytene, while in parallel late pachytene cells enter diplotene and progress through the two meiotic divisions.

### Synapsis in *Clytia* is Spo11 dependent

We identified a single conserved *Clytia Spo11* gene using genome and transcriptome data (Supplementary Figure S2) (22). We also identified other key meiosis synaptonemal complex components such as Sycp1 and Sycp3, and the repair pathway proteins Rad51, Mlh1, Dmc1 (Supplementary Figure S3, Supplementary data S3-S9). SC components are highly similar between cnidarians and many bilaterians including mammals, while more divergence is observed amongst SC components from ecdysozoan species in particular, including Drosophila and Caenorhabditis (23) (Supplementary Figure S3). We mutated the *Clytia Spo11* gene by injection into eggs of single CRISPR-Cas9 guide RNAs targeting either exon 1 or exon 2 (Figure 2A, Supplementary Figure S4). This generated F0 male and female *Spo11* mutant polyp colonies each containing small sets of insertion and deletion events around the targeted site, but no detectable wild-type *Spo11* sequence (Table 1, Supplementary Figure S4). This typical outcome of CRISPR-Cas9 mutagenesis in *Clytia* reflects very active but inaccurate Microhomology Mediated End-Joining (MMEJ) repair of DNA double strand breaks in the embryo, followed by reduction in genotypic complexity during larval metamorphosis (17). As expected, mutant alleles were invariant among jellyfish produced from the same mutant polyp colony.

**Figure 2:**
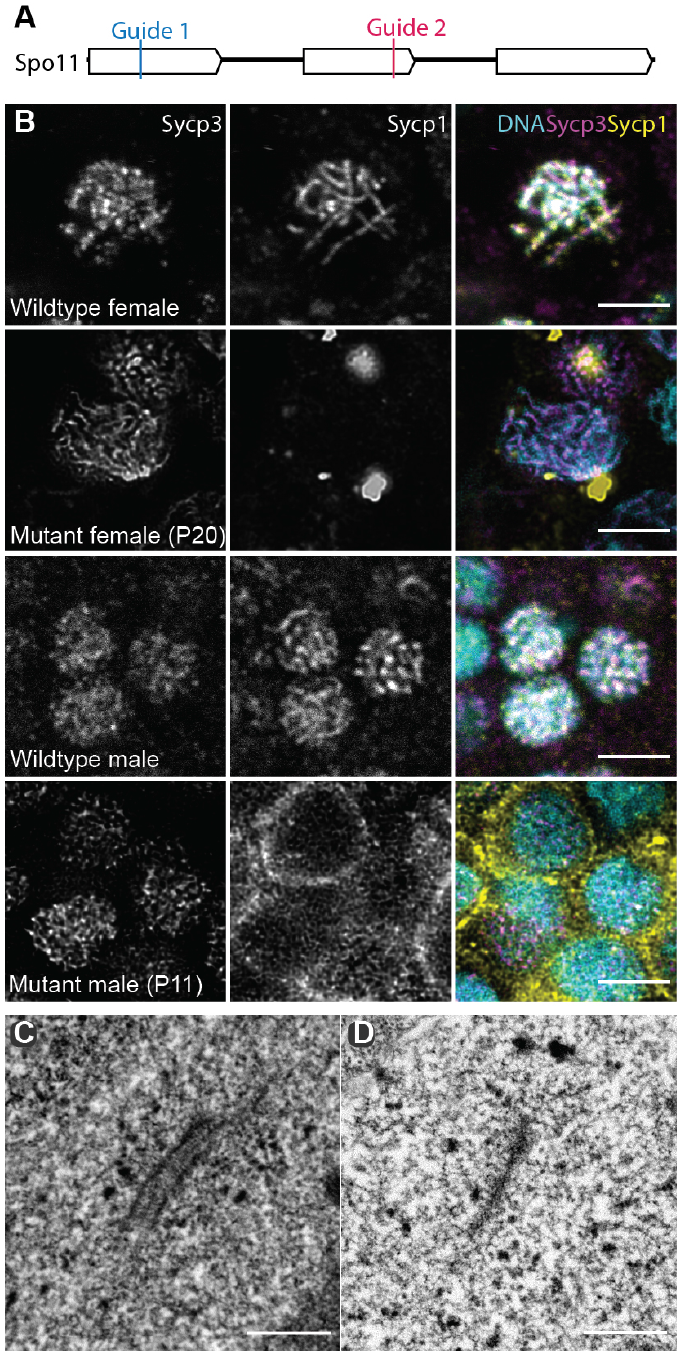
Spo11 is required for synapsis in *Clytia hemisphaerica* males and females. A. Diagram of the Spo11 exons indicating the target sites of CRISPR/Cas9 guides. B. Confocal planes of wildtype and Spo11 mutant pachytene and pachytene-like nuclei stained with anti-Sycp1, anti-Sycp3, and Hoechst. Scale bar 5 µm. C. Transmission electron microscope (TEM) image of a wildtype synaptonemal complex. Scale bar 0.5 µm. D. TEM image of a Spo11 mutant (P16) axial element. Counts in Table S1. Scale bar 0.5 µm.

**Table 1:**
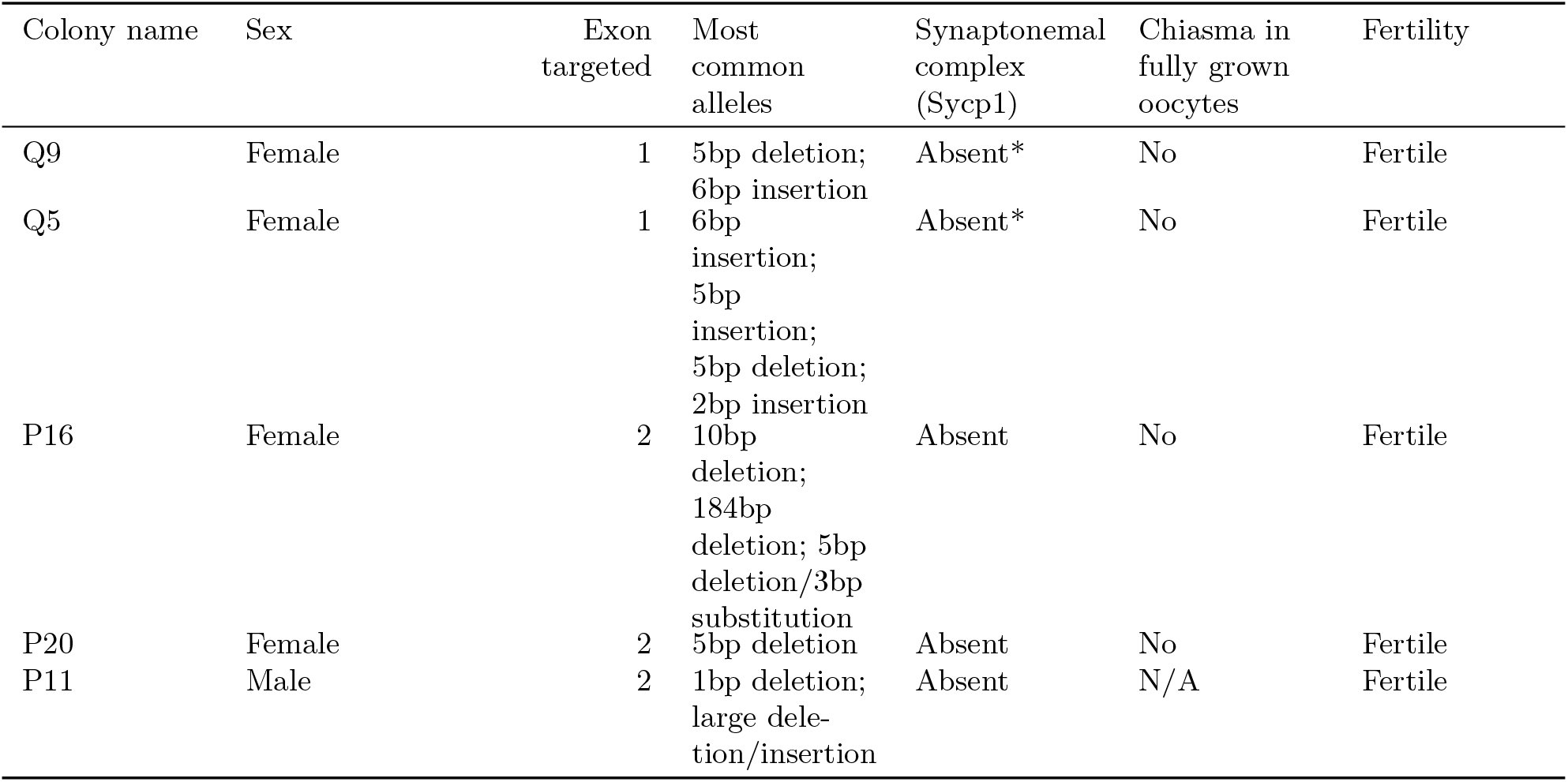
Summary of genotypes and phenotypes of Spo11 mutant colonies in this study. Asterisk indicates occasional partial SCs detected

We performed telomere FISH for two female *Spo11* mutants and could distinguish telomere bouquets characterizing leptotene/zygotene (Supplementary Figure S5), indicating that interchromosomal interactions during these early stages are not dependent on Spo11 function. Pachytene is characterized by a fully formed synaptonemal complex (SC), which mediates tight association between each pair of duplicated chromosomes (3). We visualized the SC during pachytene in both males and females using antibodies generated to *Clytia* Sycp1, which forms the transverse filaments of the SC, and Sycp3, a component of the axial elements associated with each individual chromatid pair (Figure 2B). In wildtype meiotic cells, Sycp3 is deposited along chromosome axes during zygotene, where it remains until diplotene, meanwhile Sycp1 is present along chromosomes in short stretches during late zygotene, and along the full length of chromosomes during pachytene – indicating that chromosomes are fully synapsed at the pachytene stage. In *Spo11* mutants we detected morphologically distinct pachytene-like stages with Sycp3 present along chromosome arms within both male and female gonads but Sycp1 and thus SCs were absent in the majority of pachytene nuclei, in rare cases anti-Sycp1 decorating only occasional short chromosome stretches (Figure 2B). Consistently, ladder-like synaptonemal complexes could be detected by electron microscopy in wild-type pachytene oocytes, (Figure 2C), but not in *Spo11* mutant pachytene oocytes (Figure 2D; Supplementary Table S1). The earliest observable defect in *Clytia Spo11* mutants is thus a failure in synapsis, i.e. in the correct association of paired chromosomes via SC formation.

### Chiasmata-mediated chromosome segregation fails in *Spo11* mutants

We predicted that in the absence of the SC, no crossover events would occur in *Spo11* mutants. We verified this in female gonads using an antibody raised to *Clytia* Mlh1 (Supplementary data S4), a protein that localizes to and stabilizes meiotic crossovers across many species (24, 25). In wildtype diplotene cells, clear Mlh1 foci were observed (Figure 3A top panel), with a mean of 32.8 ± 16.4 foci per nucleus (n= 21; raw counts in supplementary data S1). In contrast, diplotene nuclei in *Spo11* mutants were completely devoid of Mlh1 foci (Figure 3A bottom panel). A convenient feature of *Clytia* oogenesis for analyzing meiotic recombination events is that paired chromosomes connected at chiasmata sites can be readily observed towards the end of the oocyte growth phase. Within the enlarged oocyte nucleus (‘Germinal Vesicle’) of wild type oocytes, 15 pairs of chromosomes with 1-2 crossovers per pair are clearly distinguishable (Figure 3B top panel). In all female *Spo11* mutants, however, we consistently distinguished 30 scattered univalents at this stage (Figure 3B bottom panel), with only occasional hints of connections between them. This phenotype strongly suggests that in *Clytia* Spo11-mediated DNA double strand breaks are necessary for the formation of chiasmata, which in turn are required to maintain chromosome pairing during oocyte growth.

**Figure 3:**
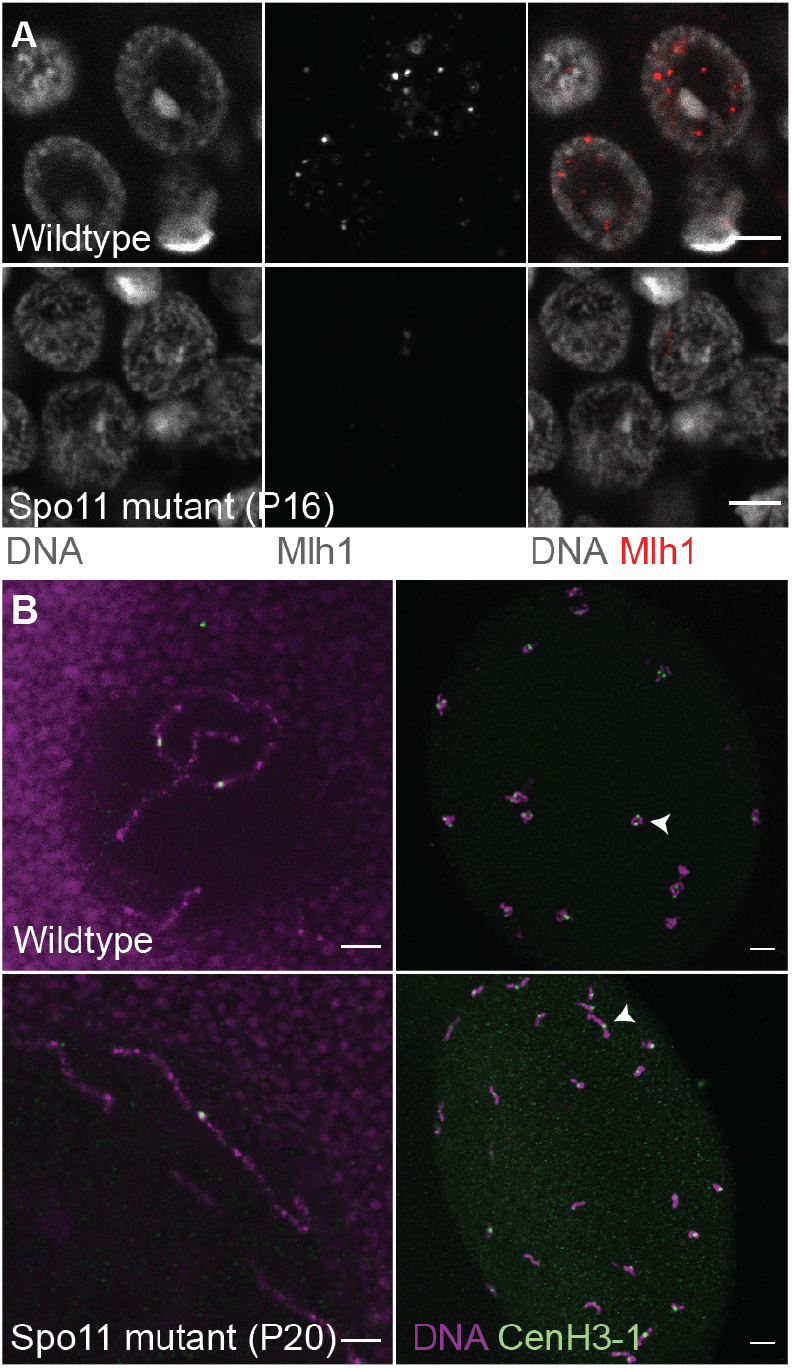
Spo11 is required for crossovers and recombination in *Clytia hemisphaerica*. A. Wildtype and *Spo11* mutant early diplotene nuclei stained with anti-Mlh1 (red in overlay). Scale bar 5 µm. B. Fully grown oocytes stained with Hoechst dye (purple) and anti-CenH3-1 (green). Top left: close up of a wildtype oocyte nucleus showing two homologous chromosomes with one chiasma. Top right: maximum intensity projection from a confocal z stack of a wildtype oocyte nucleus showing 15 pairs of chromosomes with one or two chiasma per pair. White arrow points to a bivalent. Bottom left: close up of a mutant oocyte nucleus showing 3 univalents. Bottom right: maximum intensity projection of a mutant oocyte nucleus showing 30 univalents. White arrow points to a univalent. Scale bar 5 µm.

The loss of chiasma and resulting scattering of individual duplicated chromosomes in fully grown oocytes had a severe impact on the subsequent meiotic divisions. When *Spo11* mutant oocytes are induced by MIH treatment to undergo meiotic maturation, univalents fail to align on a metaphase plate at the first division and remain distributed across defective spindle structures (Figure 4A,B first and third column). Moreover, polar body formation in *Spo11* mutant oocytes frequently aborts during anaphase I (Figure 4A,B second column). Despite these severe abnormalities in the meiotic divisions, resulting typically in retention of two or four haploid sets of chromosomes in the oocyte, *Spo11* mutant oocytes could be successfully fertilized with wildtype sperm at near wildtype frequencies. In most cases the resulting zygotes developed into planula larvae and could undergo metamorphosis to the polyp stage (Supplementary Table S2). Similarly, *Spo11* mutant males show no loss of fertility, producing functional sperm capable of fertilizing wildtype oocytes to generate metamorphosis-competent larvae (Supplementary Table S2). These *Spo11* mutant x wildtype progeny could provide a fascinating opportunity to address mechanisms of aneuploidy management.

**Figure 4:**
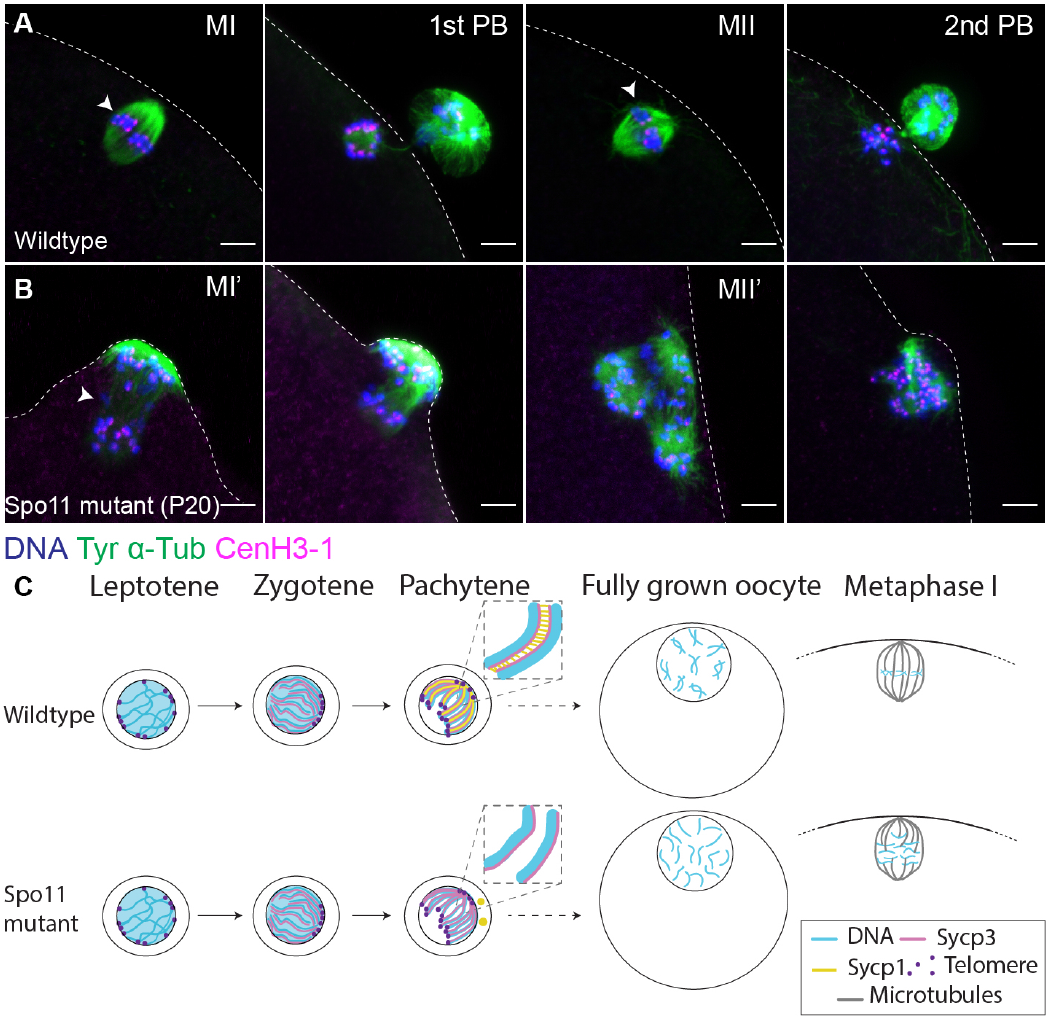
Meiotic division defects in *Spo11* mutant oocytes. DNA is stained with Hoechst dye (blue), microtubules are stained with antityrosinated tubulin (green), and centromeres are stained with antiCenH3-1 (magenta). Dotted lines contour the oocyte surface. A. Wildtype meiotic divisions. From left to right: metaphase I (MI); first polar body (1st PB) emission; metaphase II (MII); second polar body (2nd PB). White arrows indicate metaphase plates; Scale bar 5 µm. B. Spo11 mutant meiotic divisions. From left to right: metaphase I (MI’), white arrow indicates where the metaphase plate should be; failed emission of the first polar body; metaphase II (MII’); Spo11 mutant oocyte at the time of second polar body emission. Scale bar 5 µm. C. Summary of meiotic progression from leptotene to the first meiotic division in wildtype and Spo11 mutants. DNA (blue), Sycp3 (magenta/pink), Sycp1 (yellow), telomeres (purple) and microtubules (grey).

In this study we show that meiosis in *Clytia hemisphaerica* relies on classical mechanisms of homologous chromosome synapsis and the resulting chiasma are required to maintain homologous chromosome pairs during oocyte growth (Figure 4C). The sequence of meiotic events and their interdependencies revealed by *Spo11* knockout in *Clytia* closely match those well described across eukaryotic model organisms including mice, zebrafish, budding yeast, filamentous fungi and plants including *Arabidopsis thaliana* (1). They are thus likely governed by ancestral and conserved mechanisms. Within animals, however, not all species share the same dependencies between Spo11 function and subsequent events. Alternative modes of DSB-independent synapsis and pairing have notably been described in the traditional animal model species *C. elegans* and *D. melanogaster* (6, 7). In *D. melanogaster*, male chromosomes show no recombination, while chromosome pairing in females initiates prior to meiotic entry in proliferating germ cells (26, 27). This “pre-pairing” mode may represent a specialization to facilitate later SC formation within the context of developing germ cells within the germarium. In *C. elegans*, recombination-independent chromosome pairing occurs at “pairing centers”, specialized sites at chromosome ends, which also promote initiation of synapsis (28). Involvement of pairing centers in synapsis likely evolved within the *Caenorhabditis* clade, since a classic role for Spo11 in chromosome pairing, synapsis, and crossover formation has been uncovered in another nematode species, *Pristionchus pacificus* (14). In the planarian *Schmidtea mediterranea*, pairing and synapsis in males are not dependent on Spo11, and it is proposed that the telomere cluster plays a role in initiating SC formation, which in turn drives homolog pairing (12). SC nucleation mechanisms are as yet unknown in this species.

Functional studies in phylogenetically distinct species provide invaluable insights into both conserved or divergent meiotic mechanisms. Along with practical advantages including transparent, simply organized gonads that are accessible to manipulation and observation, the jellyfish *Clytia hemisphaerica* offers a valuable evolutionary comparative perspective on meiotic mechanisms, which in this model are largely conserved from a distant eukaryotic ancestor.

## Materials and Methods

### Clytia hemisphaerica cultures and maintenance

*Clytia hemisphaerica* cultures were maintained as described previously (29). Several wildtype strains were used in this study, with consistent results, including female strains Z11, Z4B, A2, Z21, Z28; and male strains Z13, Z23, A3. CRISPR-Cas9 mutant jellyfish were generated by injecting Z11 oocytes and fertilizing them with Z13 sperm.

### Gene identification and homology

*Clytia hemisphaerica* genes, for antibody, guide RNA, and FISH probe design, were identified via BLAST against the Clytia genome http://marimba.obs-vlfr.fr/ (22). Gene alignments were made using MAFFT v7.453 L-INS-i (30), and gene phylogenies generated with IQ-TREE(31) using a LG+F+R4 model. Gene trees and alignments are available as supplementary data S3-S9.

### Spo11 CRISPR-Cas9 knockouts

Spo11 mutants were generated as described previously(17, 19). Candidate guide RNAs with predicted cut sites between microhomologies were identified using http://www.rgenome.net/mich-calculator/, and offtarget sites using http://crispor.tefor.net/. crRNA sequences are listed in table S3. Spo11 small guide RNA (sgRNA) was generated by hybridizing 200 µM crRNA and 200 µM tracrRNA in the presence of 1x Hybridization buffer (Integrated DNA Technologies, Coralville, IA), 95°C for 5 min, cooled to 25°C at - 0.1°C/sec. Two separate sgRNAs were injected, targeting regions of exon 1 and of exon 2. Prior to injection, 0.5µl of hybridized crRNA/tracrRNA (60µM) was mixed with 2µl Cas9 protein (10µM), incubated at room temperature for 10 minutes, adjusted with 0.84µl Cas9 Buffer (10 mM Hepes, 350 mM KCl), and centrifuged at 14,000 rpm for 10 minutes at 12°C. Injection, fertilization and subsequent metamorphosis of larvae was conducted as described previously (17, 19).

### Genotyping

Genomic DNA was extracted from a single jellyfish using DNeasy blood/tissue extraction kit (Qiagen). DNA around the target site was amplified using Phusion DNA polymerase (NEB, Ipswich, MA). PCR primer sequences are listed in table S3. PCR products were cleaned up and sequenced. Genotypes of Clytia mutant strains are in Table 1 and Supplementary Figure S4.

### Antibodies

Antibodies recognizing Clytia Sycp1 were generated in rabbits using two peptides IRNWKSEKEMELKMKD-Cys (75-90aa) and Cys-PKAMTPKTPNMRYS (833-846aa). Antibodies recognizing Clytia Sycp3 were generated in rats using two peptides ENAPAEEAPAISGK-Cys (25-38aa) and GRKRPAPHISHT-Cys (39-50aa). Antibodies recognising Clytia Mlh1 and CenH3-1 were generated in mice using a recombinant protein generated in E. coli against the full length optimized sequence. Antibodies recognising Clytia Piwi1 were raised in rabbits using a recombinant protein generated in E. coli against SGEPVQILTNYFKVDKMPRFEGLHQYVVAFDPDIQSQKLKGFLLFSMQDVIGEVKVFDGM-SLFLPRKLAEPVVERCVETRDGSSIKVKITHTNEVPVNSPQVVQLM (115-220aa) fused with an N-his Tag. All antibodies were generated and affinity-purified by Proteogenix (Schiltigheim, France).

For immunostaining, anti-Sycp1 antibodies were diluted 1:2000 for sycp1#4 and 1:1000 for sycp1#2. Sycp3#2 was diluted 1:400. Mlh1#2 was diluted 1:250. Piwi#1 and Piwi#2 were diluted 1:2000. CenH3-1#2 was diluted 1:1000. Secondary antibodies (anti-mouse Mlh1; anti-rabbit Sycp1, Piwi) were diluted 1:200. Rat monoclonal anti tyrosinated tubulin YL1/2 was diluted 1:500 (Thermo Fisher Scientific).

### Immunostaining

For anti-Sycp1, anti-Sycp3 and anti-Mlh1 staining, samples (whole 1 week jellyfish or dissected gonads) were fixed in 1% formaldehyde in methanol at -20°C for at least 2 hours (up to overnight). Methanol fixed samples are left to warm gradually to room temperature (∼30 mins) before rehydration to PBS. For anti-tubulin, anti-CenH3-1, and anti-Piwi1 staining, samples were fixed for 2h at room temperature in IF fix (Hepes pH 6.9, 0.1M, EGTA pH 7.2 50 mM, MgSO4 10mM, Maltose 80 mM, Triton 100 × 0.2%, 4% paraformaldehyde) for 2 hours before proceeding to washes and staining.

Anti-tubulin and anti-CenH3-1 staining included a methanol series: PBS-Triton 0.01% 3 × 10 mins, 50% methanol/50% PBS-Tween 0.1% 1 × 10 mins on ice, 100% methanol 2 × 10 mins on ice (or store in -20°C after 2nd wash up to a week), 50% methanol/50% PBS-Tween 0.1% 1 × 10 mins at RT. All antibody staining then proceeded with the following washes: PBS-Triton 0.01% 3 × 10 mins, PBS-Triton 0.2% 40 mins, PBS-Triton 0.01% 2 × 10 mins, PBS/BSA 3% 20 mins-1h. Primary antibodies were diluted with PBS/BSA 3% and incubated with the sample at 4°C overnight. The next day, wash PBS-Triton 0.01% 3 × 10 mins. Dilute secondary antibody in PBS-Triton 0.01% and Hoechst 33342 1:5000, incubate with sample 2h at room temperature or overnight 4°C. Wash in PBS-Triton 0.01% 4 × 5 min, mount in Citifluor antifade mountant (Citifluor-EMS).

### Telomere FISH

Telomere FISH was conducted using Cy3 and Alexa647-labeled G-Rich telomere probe (Eurogentec, PN-TG020-005, PN-TG050-005) targeting repeats of TTAGGG. Probes were resuspended in formamide at 50 M. A working aliquot was stored at 4°C for regular use, remaining aliquots were stored at -20 for longer-term storage. Hybridization buffer solution was made fresh each time before starting the FISH protocol: 20mM Sodium phosphate Na2HPO4 pH 7.4, 20mM Tris-HCl pH 7.4, 60% Deionized Formamide, 2xSSC, tRNA 1x, Heparin 1x.

FISH was conducted as follows: samples (up to 1 week jellyfish (5-8 per tube), isolated oocytes) were fixed overnight at room temperature in HEM fix (Hepes pH 6.9, 0.1M, EGTA pH 7.2 50 mM, MgSO4 10mM, 4% formaldehyde), washed 3x 10 mins with PBS-Tween 0.1%, and then dehydrated and rehydrated in methanol on ice (PBST/methanol 50% 1x 10 min, Methanol 100% 2 × 10 min, PBST/methanol 50% 1 × 10 min). Samples were then washed in 2xSSC-0.1% Tween (pH 7), 3x 5min. They were incubated in RNaseA solution (100mg/ml) in an oven at 37°C for 1 h. Then washed 3x 5 mins in 2xSSC-0.1% Tween (pH 7) at room temperature. Then transferred to 50% hybridization buffer/2xSSC at 80°C for 5 mins in a water bath. In parallel, the probe (1µl in 99µl hybridization buffer) was heated 90°C for 5 mins. As quickly as possible, we removed as much 50% hybridization buffer/2xSSC as possible and added the probe to the sample, gently flicking the tube to mix. The sample was then incubated 10 mins at 85°C in a water bath. Subsequently, samples were placed in the dark at room temperature for 1 hour for hybridization. After hybridization, samples were washed 2x 10 mins with 2xSSC-0.1% Tween at 60°C in an oven (2xSSC-0.1% was preheated to 60°C prior to washing). Samples were washed once with 2xSSC-0.1% Tween at room temperature, then stained with Hoechst dye 33342 1:5000 (30 mins - 1h), washed 3×5 mins with 2xSSC, and mounted in Citifluor.

### In-situ Hybridization

In-situ hybridization probes were produced as described previously (32), and in-situs performed following the urea-based protocol (33).

### Transmission Electron Microscopy (TEM)

For TEM, gonads were dissected from 4 and 5 day old female jellyfish in 400 µM menthol in seawater (Sigma-Aldrich, #M2772, diluted from a 1M stock solution in ethanol). Fixation and embedding in Epoxy resin were performed as described previously (34).

### Fertilization Experiments

Embryos were generated as described previously (29) with the following crosses: mutant female x wildtype male, mutant male x wildtype female, with a control of wildtype female x wildtype male. Mutant or wildtype eggs were collected and added to glass dishes with the same volume of liquid. For female mutant crosses, the same volume of wildtype sperm was added to mutants as to wildtype. If wildtype fertilization rate was low or development disrupted, indicative of too much or too little sperm, results were discarded. In all experiments, adequate sperm concentration was verified by the presence of ∼12 sperm around each egg. Raw counts are available in supplementary file 2.

### Meiotic maturation

Meiotic maturation was induced as described previously (19). Briefly, gonads were dissected from mutant and wildtype female jellyfish adapted to an afternoon light cycle, and were maintained overnight in dishes with the light on. The following morning, fully grown oocytes were isolated, and the MIH peptide (WPRPamide) was added to seawater containing the oocytes for a final concentration of 100 nM. Oocytes were fixed every 5 minutes, starting 25 minutes after addition of MIH.

## Acknowledgements

We thank Tsuyoshi Momose for advice and help with CRISPR/Cas9 protocols and culture of mutant colonies; Lisa Rouressol for assisting with exploration of male Clytia gametogenesis; Régis Lasbleiz and Axel Duchene at the Marine Resources Centre (CRBM) and Sébastien Schaub at the imaging platform (PIV) of Institut de la Mer de Villefranche. We thank the Huynh and Houliston labs for helpful comments on the manuscript. Transmission Electron Microscopy was performed at the Plateforme Commune de Microscopie Electronique, Université Côte d’Azur. Funding: CRBM and PIV are supported by EMBRC-France, whose French state funds are managed by the ANR within the Investments of the Future program under reference ANR-10-INBS-02. This project received funding from the European Union’s Horizon 2020 research and innovation programme under the Marie Skłodowska-Curie grant agreement No 841433 (JOLI), from the CNRS-INSB “Diversity of Biological Mechanisms” program, and from the European Union’s Horizon 2020 research and innovation programme No 730984, ASSEMBLE Plus project, JRA3. The J.-R.H. lab is supported by CNRS, INSERM, Collège de France, La Fondation pour la Recherche Médicale (FRM) (Equipes FRM DEQ20160334884), Agence Nationale de la Recherche (ANR) (ANR-15-CE13-0001-01, AbsCyStem) and the Bettencourt Schueller Foundation. Authors contributions: Conceptualization: J-RH, CM, EH; Writing: CM, JRH, EH; Investigation: CM, HC, SP; Formal analysis: J-RH, EH, CM. Competing interests: none. Data and materials availability: All data is available in the manuscript or the supplementary materials.

