## Supplementary figures and images for "Conserved meiotic mechanisms in the cnidarian *Clytia hemisphaerica* revealed by Spo11 knockout"

### CenpA_aa_extended_alignment_2.fa.contree.pdf

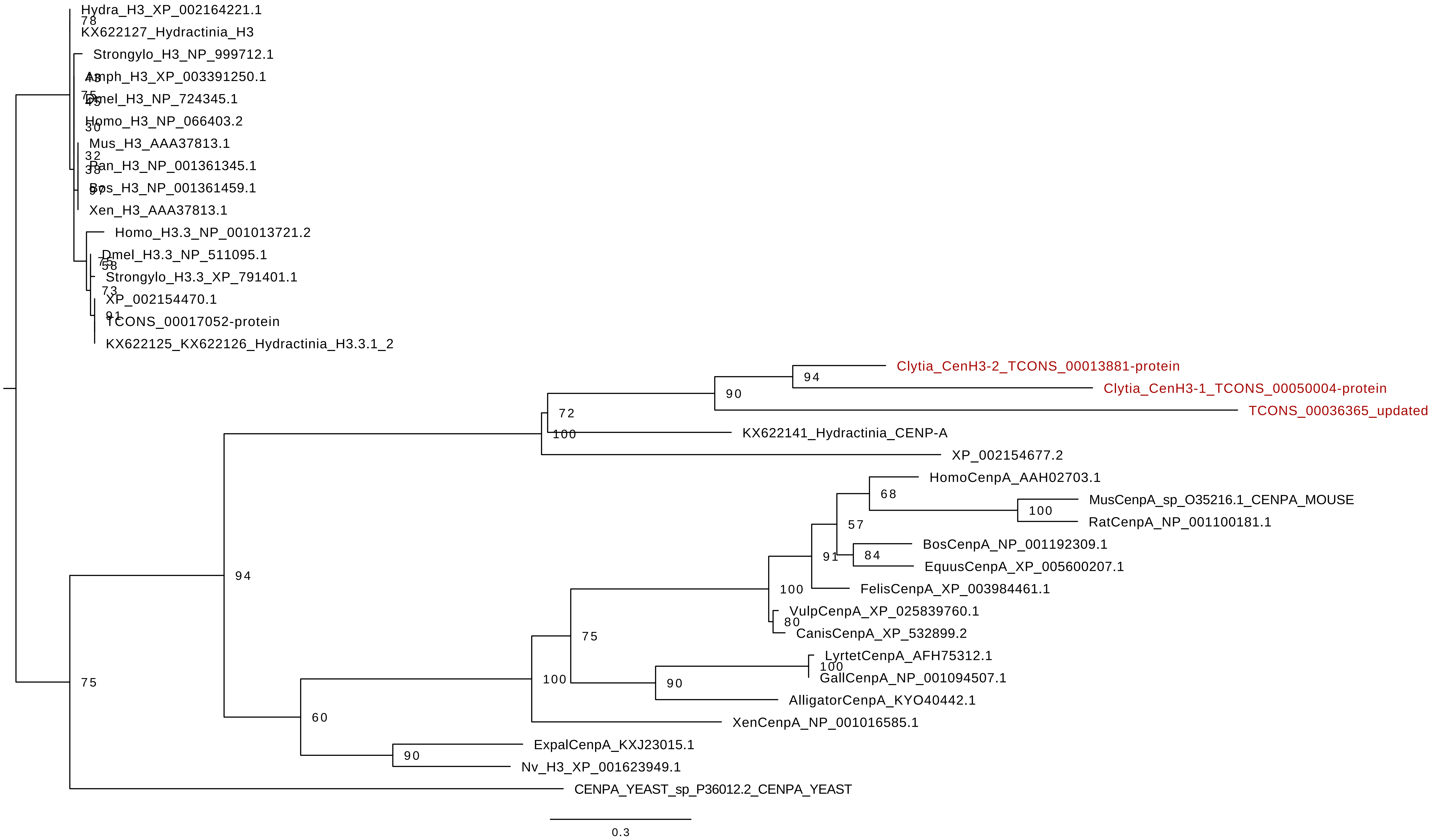

### mlh1_alignment_2.fa.contree.pdf

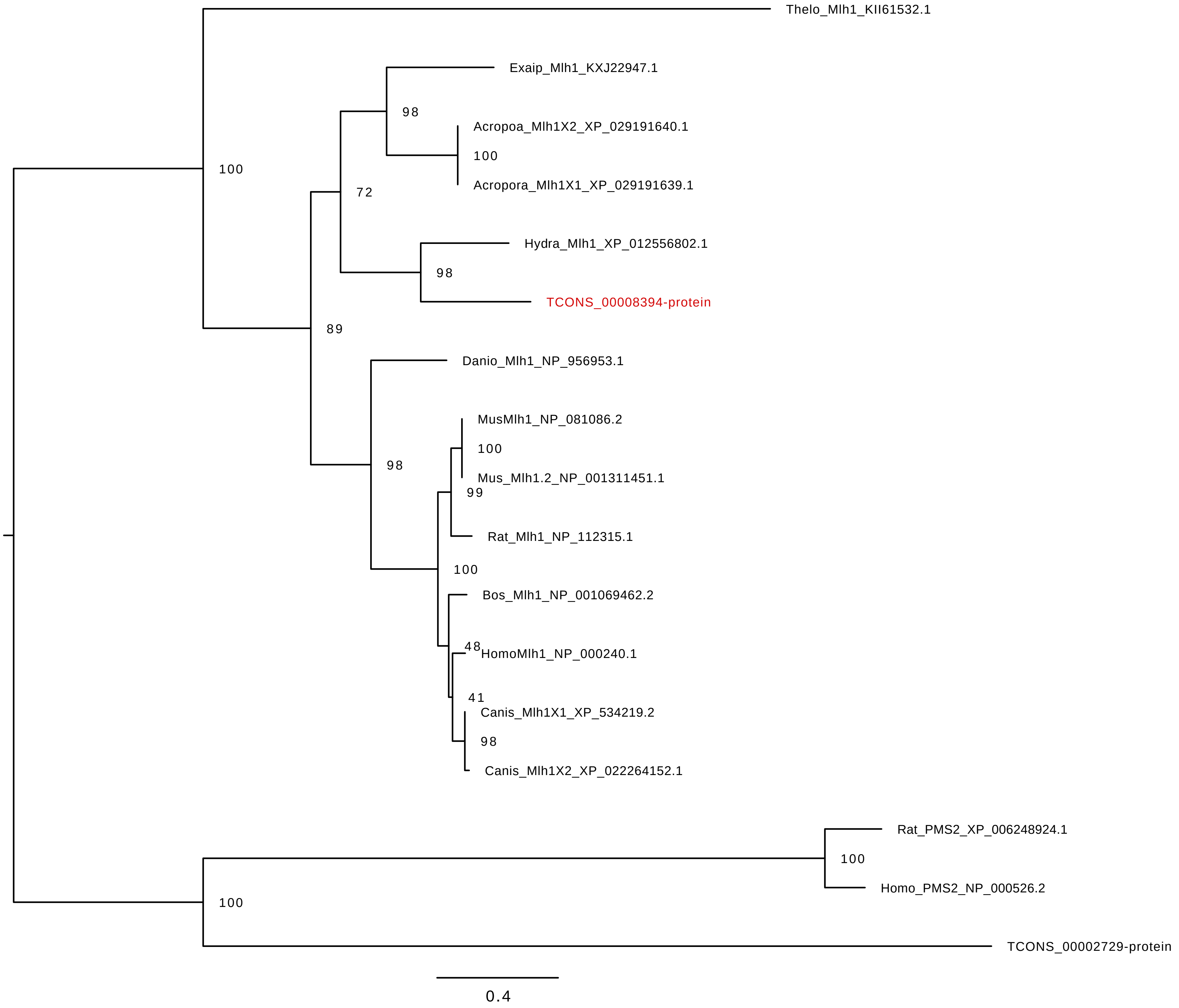

### piwi_aa_al.fa.contree.pdf

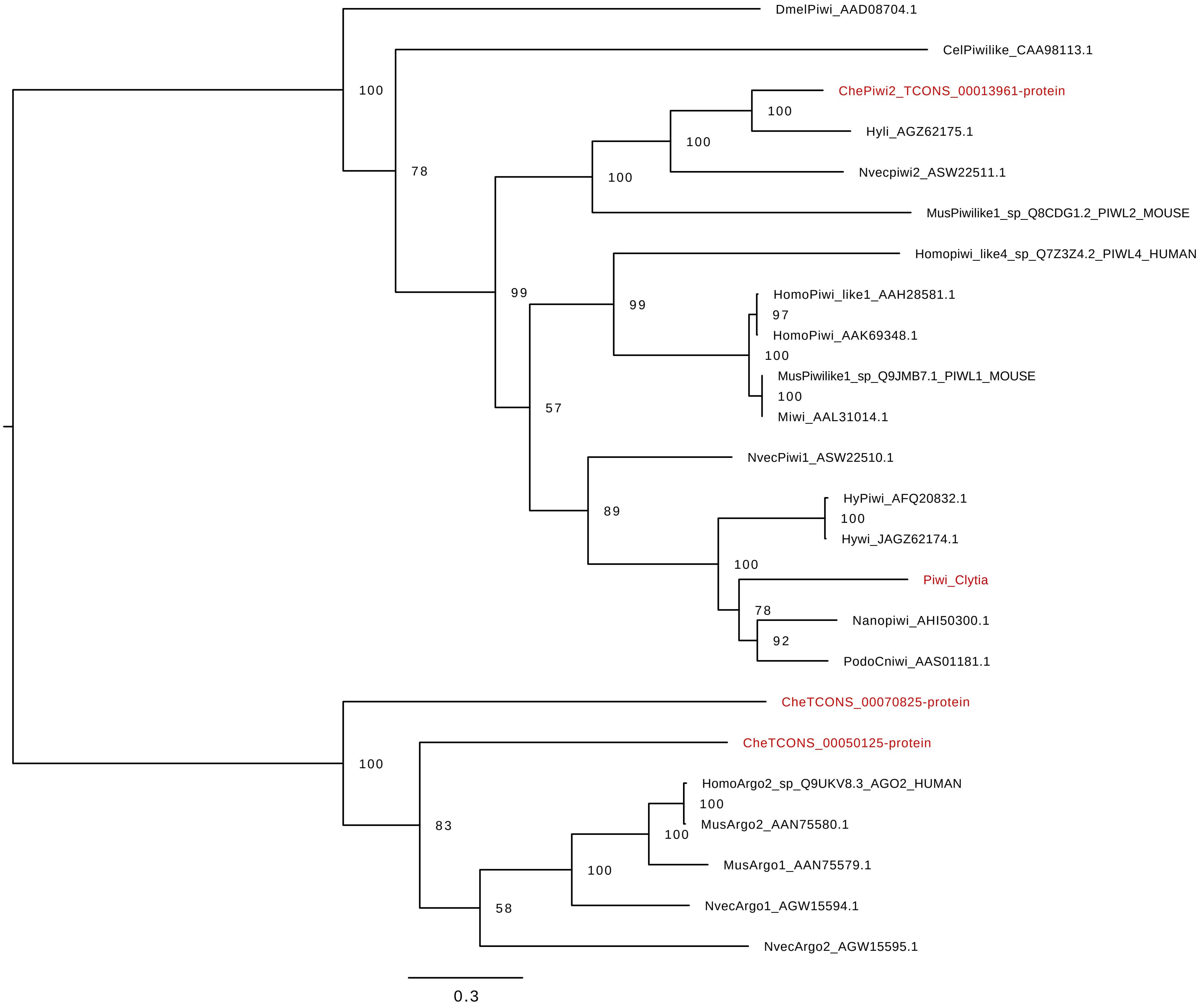

### rad51_aa.aln.contree.pdf

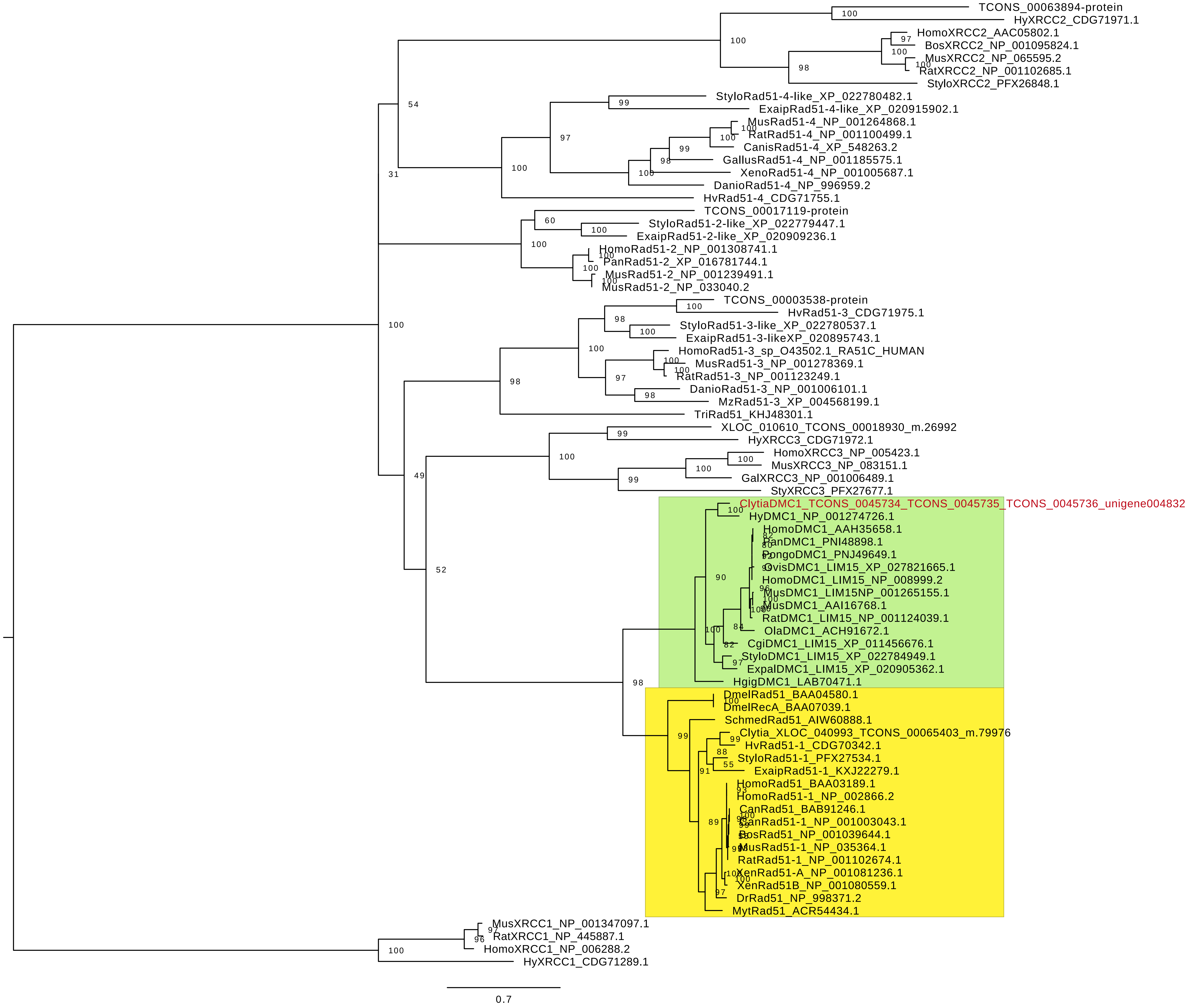

### rad51_aa.aln.contree.png

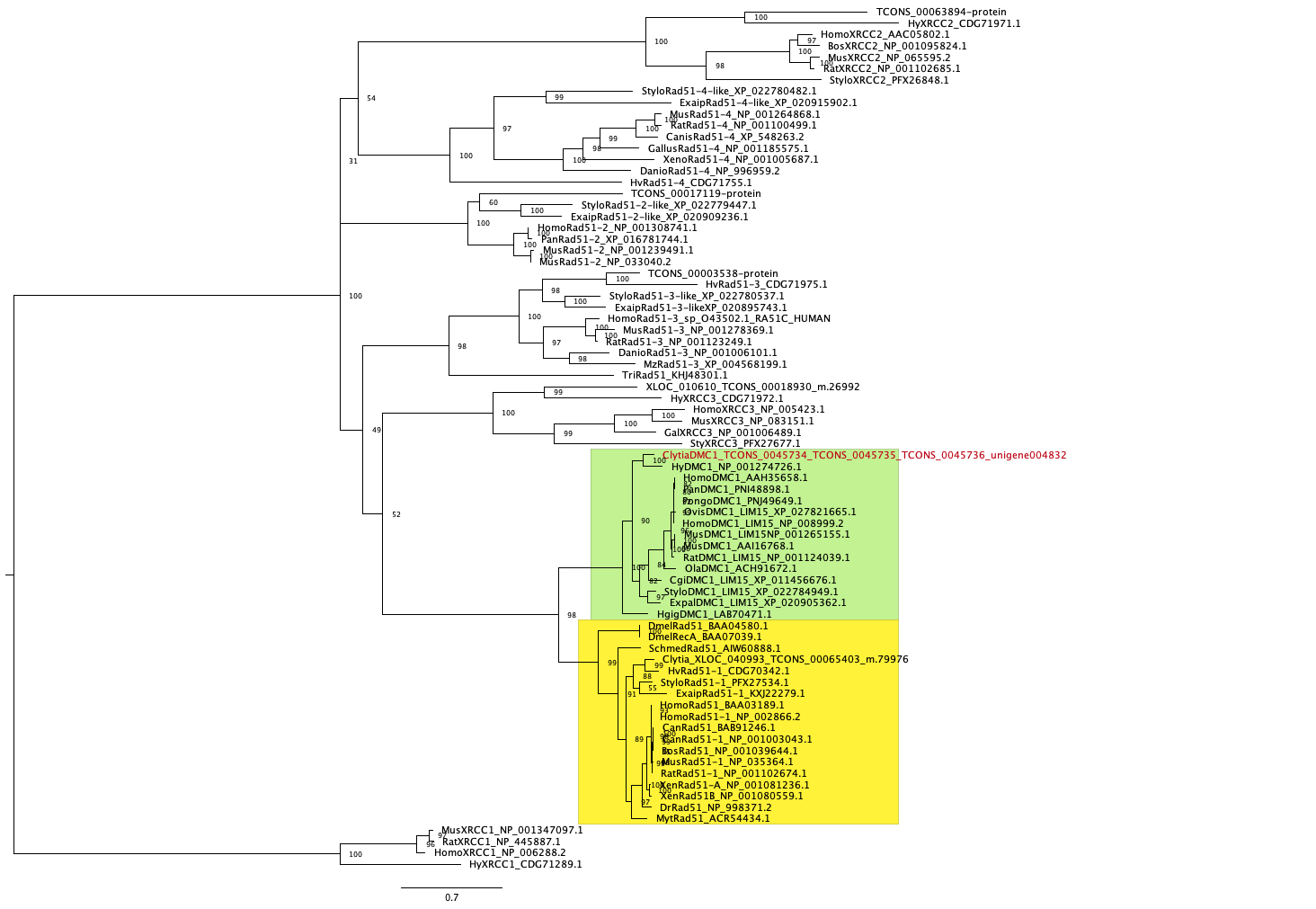

### spo11.aln.contree.pdf

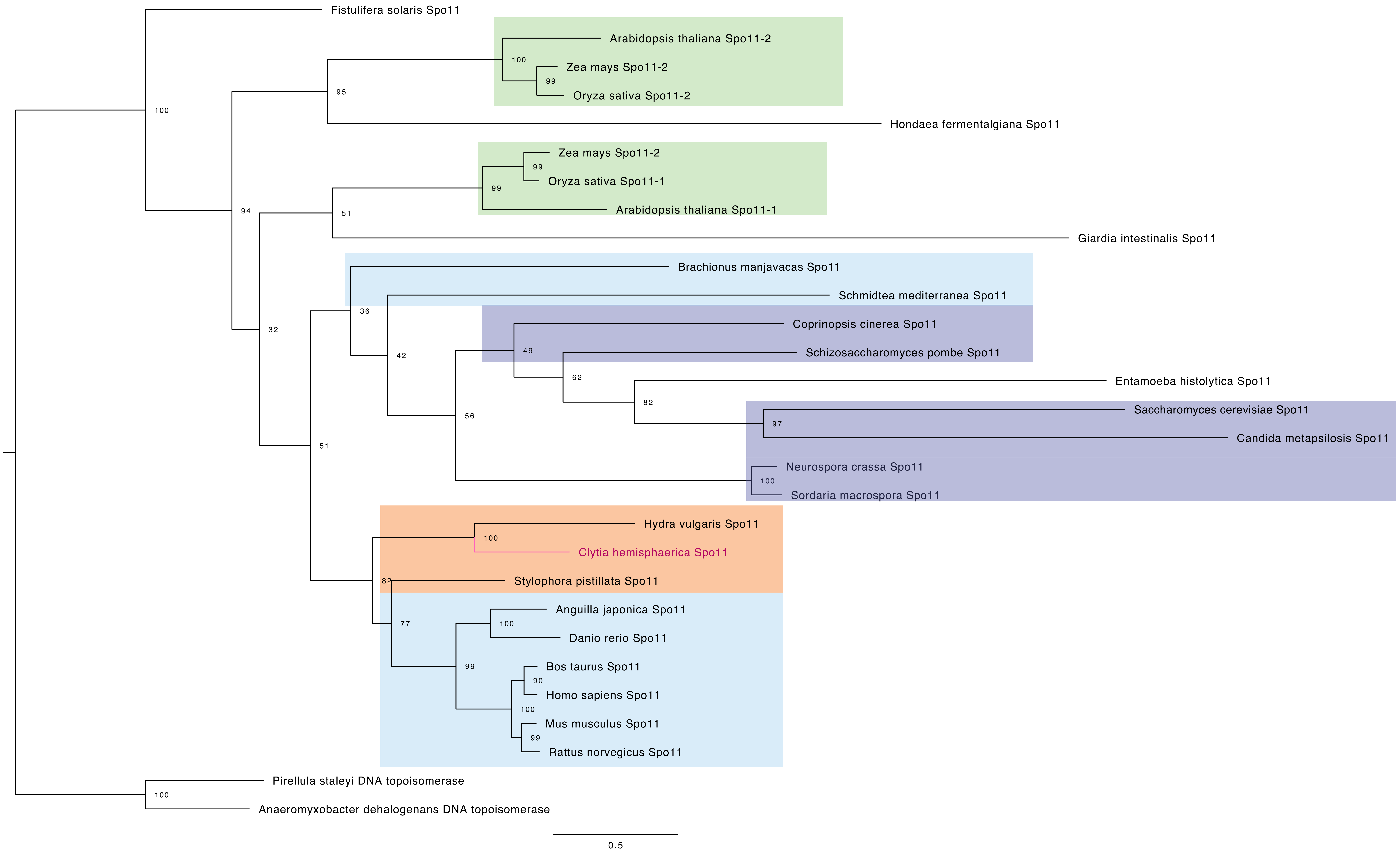

### sycp1_aa.aln.contree.pdf

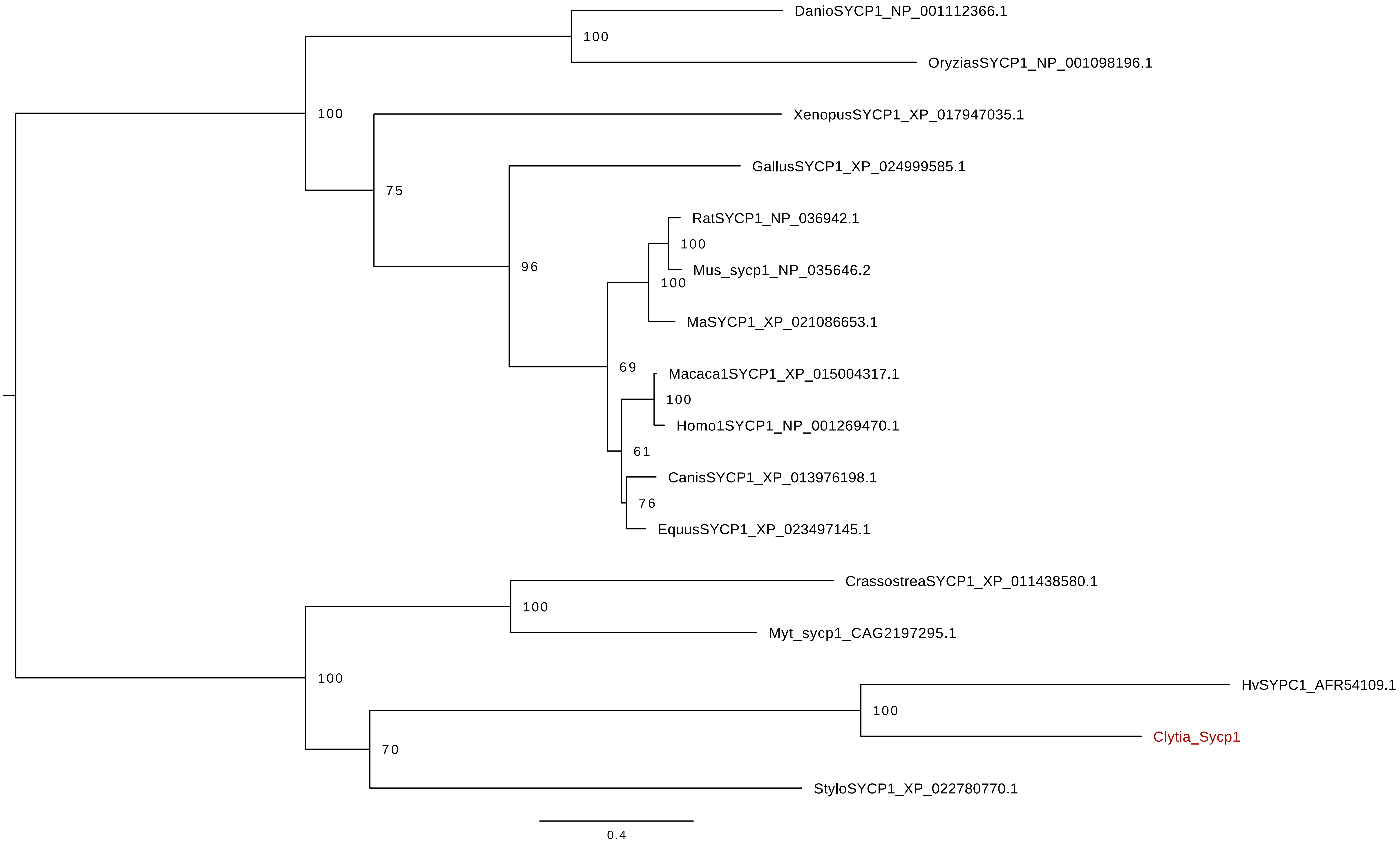

### sycp1_aa_ecdysozoa.aln.contree.pdf

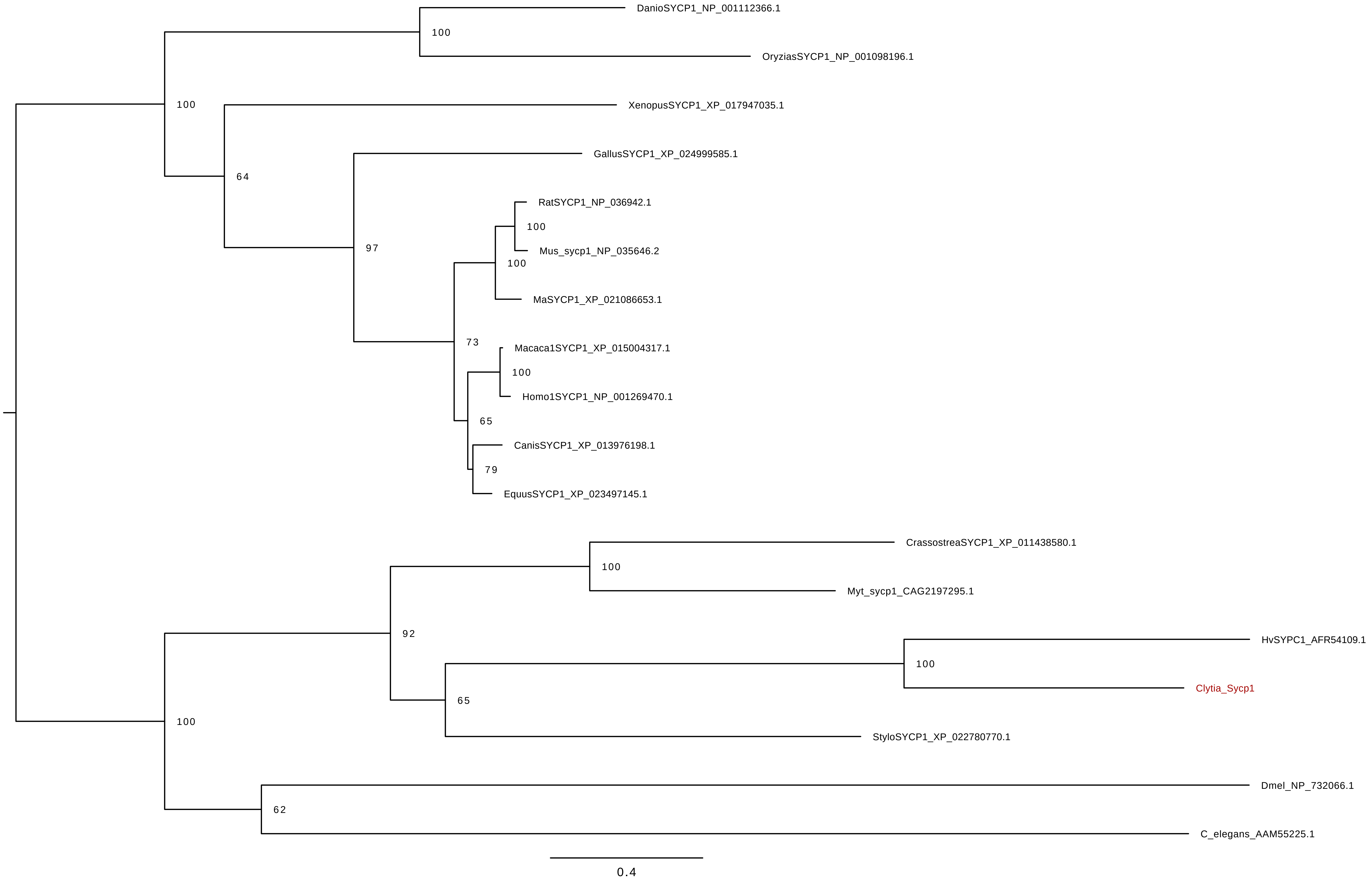

### sycp2_sycp3.aln.contree.pdf

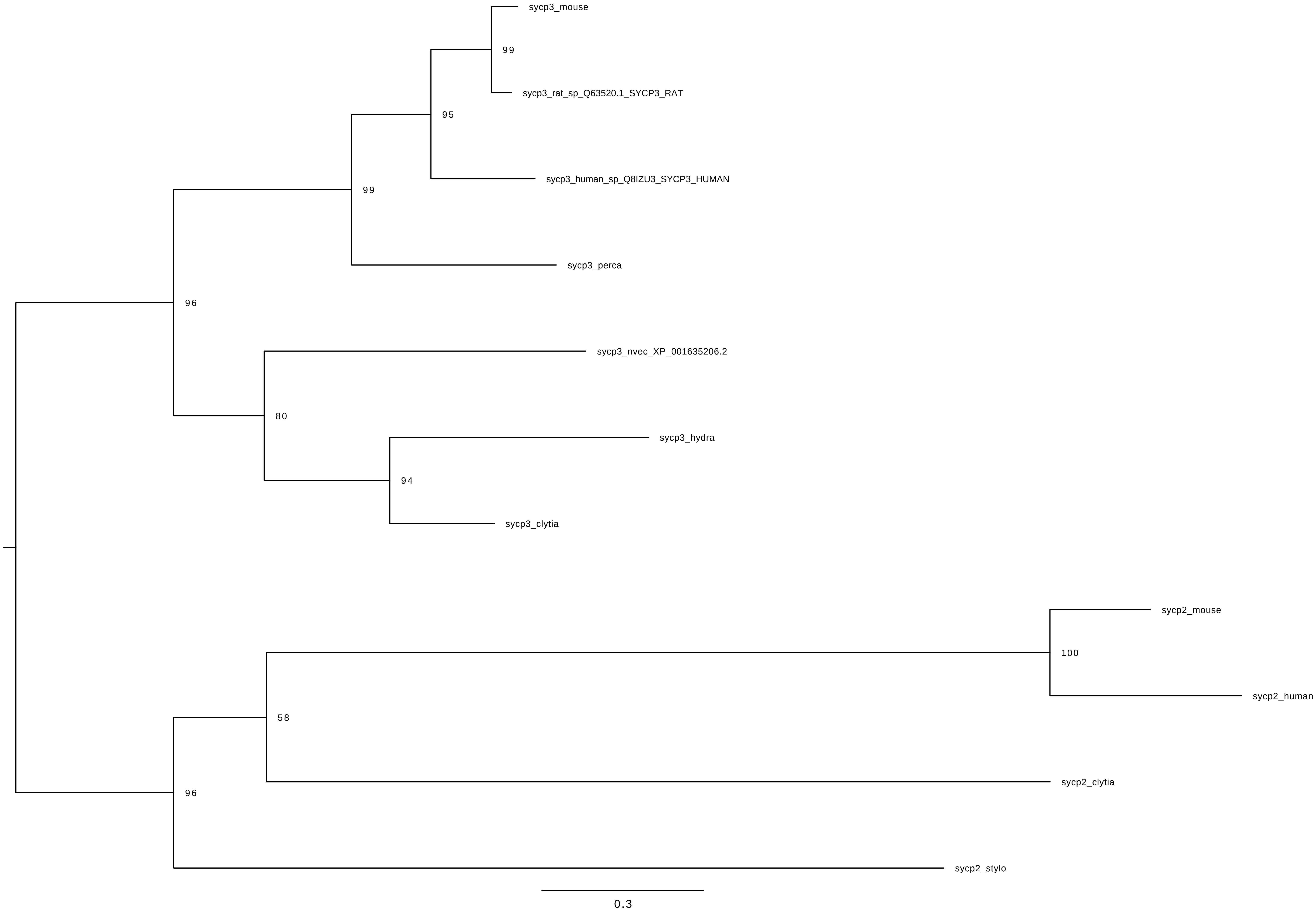
