## Supplementary figure for "Conserved meiotic mechanisms in the cnidarian *Clytia hemisphaerica* revealed by Spo11 knockout"

### Supplementary Figures and Tables

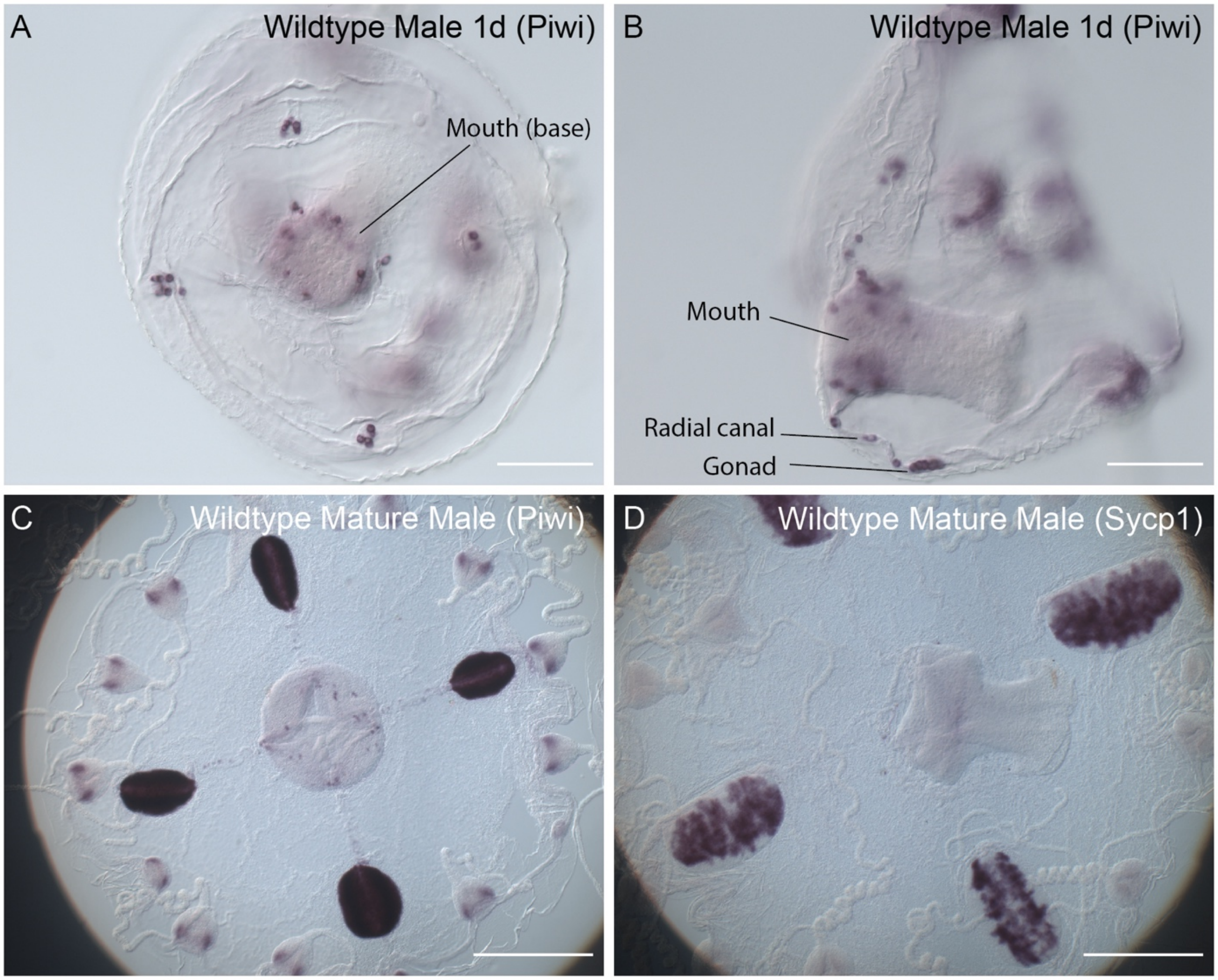


**Fig. S1.**

Expression of *Piwi1* and *Sycp1* genes in *Clytia hemisphaerica* males. **A-B.** In situ hybridisation detection of *Piwi* mRNA in 1 day male jellyfish. A - view from underside of jellyfish, showing the base of the manubrium and 4 sites of *Piwi* expression along the radial canal at sites of the future gonad. B - view from the side. **C.** *Piwi* expression in a fully sexually mature male jellyfish. **D.** *Sycp1* expression in a fully sexually mature jellyfish. Scale bars 100µm for A-B, 500 µm for C-D.

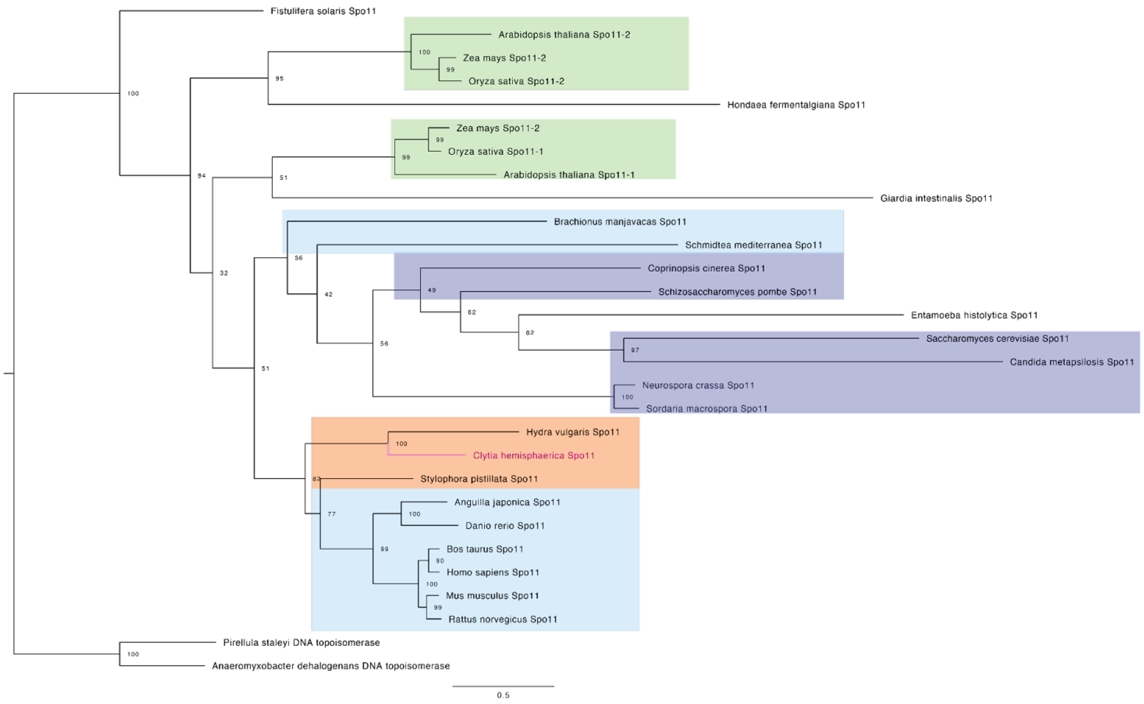


**Fig. S2.**

Maximum likelihood phylogeny of selected Spo11 protein sequences, using the LG+F+R4 model, with bacterial DNA topoisomerase proteins used as the outgroup. Bootstrap support from 1000 replicates is indicated out of 100 at each node. *Clytia hemisphaerica* Spo11 is highlighted in magenta. Blue = Bilateria, Orange = Cnidaria, Purple = Fungi, Green = Plantae. Eukaryotic Spo11 proteins (non-opisthokont and non-plant) are not coloured.

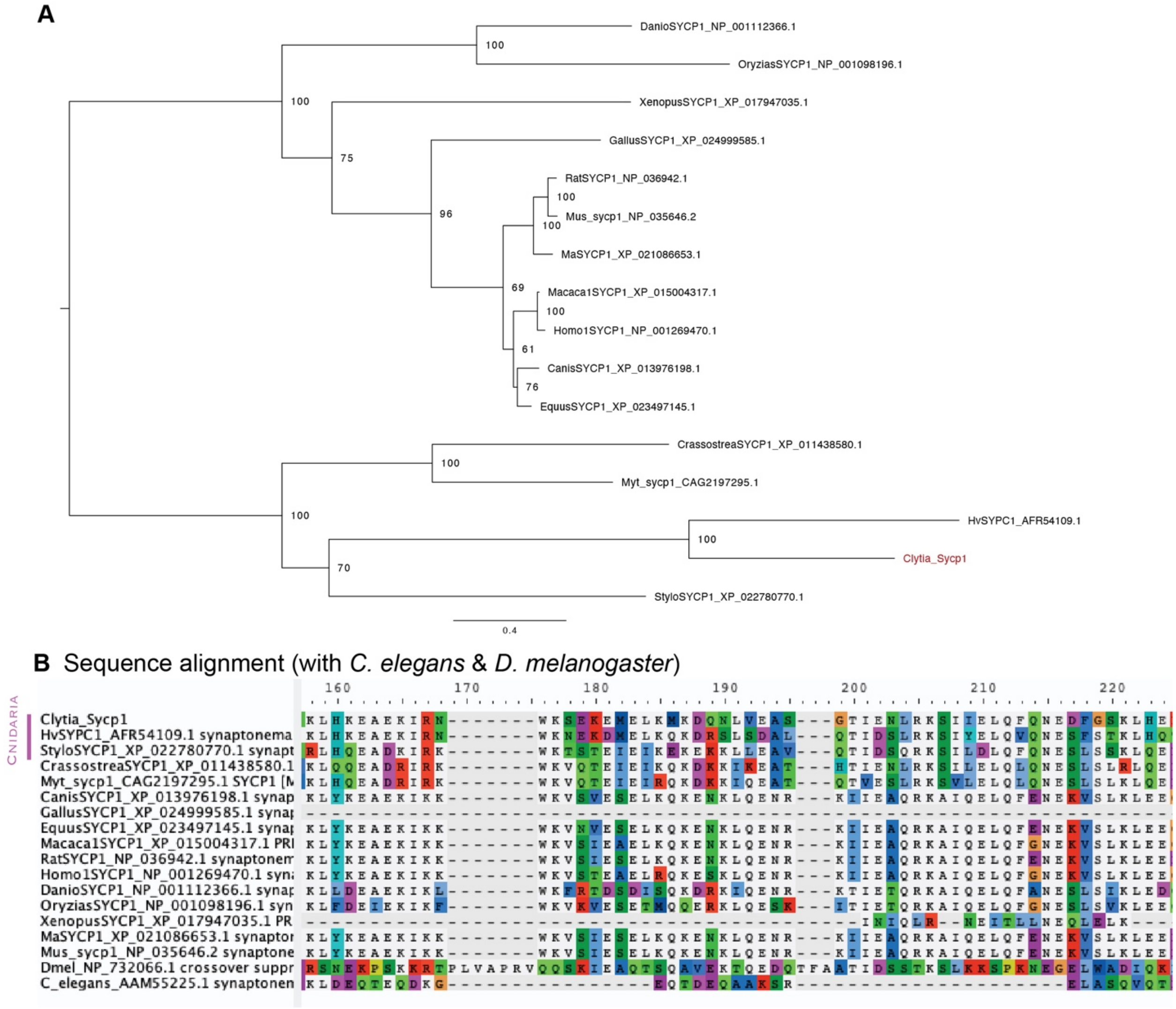


**Fig. S3.**

**A.** Maximum likelihood phylogeny of selected Sycp1 protein sequences, using the LG+F+R4 model. Bootstrap support from 1000 replicates is indicated out of 100 at each node. *Clytia hemisphaerica* Sycp1 is highlighted in magenta. **B.** Part of a multiple sequence alignment of selected Sycp1 protein sequences showing a region of sequence conservation (differences from majority rule consensus are highlighted). Sequence alignment includes divergent *Drosophila melanogaster* c(3)g and *Caenorhabditis elegans* SYP-1 sequences.

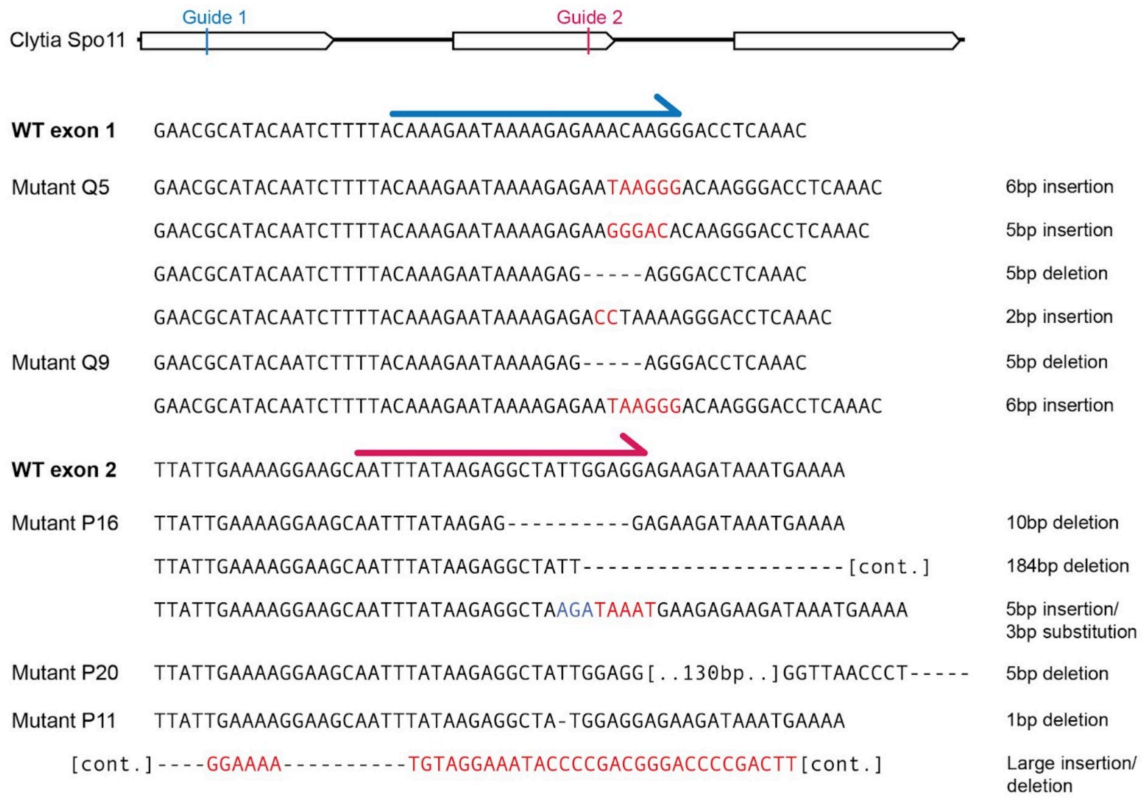


**Fig. S4.**
Sequences of most common mutations in *Spo11* exon 1 and exon 2 mutants. WT= wildtype sequence. crRNA Guide 1 (targeting exon 1) is shown as a blue arrow, crRNA Guide 2 (targeting exon 2) is shown with a magenta arrow. Insertions are highlighted as red text, substitutions as blue text, deletions as dashes.

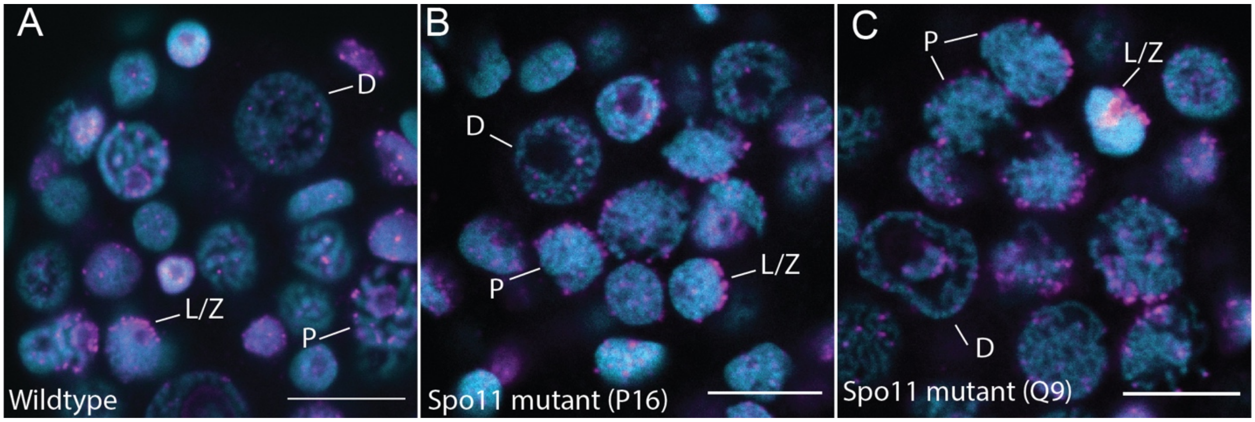


**Fig. S5.**
Confocal images showing Telomere FISH staining (magenta) overlaid on Hoechst (cyan). **A.** Wildtype female ovary. **B & C.** Spo11 mutant gonads (two different strains). L/Z - leptotene/zygotene, P - pachytene, D - diplotene. For clarity, not all nuclei are labeled. Scale bar 10 μm.

**Table S1.**

Number of distinct synaptonemal complexes (SC) detected in EM images of nuclei containing clear axial elements (AE). Counts were conducted blind to strain type.

| **Colony** | **Distinct SC / Total AE** | **Total number of nuclei examined** |
|---|---|---|
| Wildtype | 11/43 | 107 |
| Q9 | 0/20 | 46 |
| Q5 | 2/21 | 46 |
| P16 | 1/48 | 80 |

**Table S2.**

Fertility of jellyfish from mutant and wildtype colonies. Numbers of experiments are shown in parentheses. Jellyfish from each colony were crossed with wildtype females (P11) or males (all others). Successful development to the planula stage was assessed 3 days after fertilization.

| Colony | Mean percentage of eggs fertilized/ total eggs* | Mean percentage of planula/ total fertilized eggs |
|---|---|---|
| Wildtype | 93.8% (n=6) | 95.4% (n=4) |
| P11 (male) | 99.5% (n=3) | 94.0% (n=3) |
| P16 | 94.8% (n=4) | 81.9% (n=3) |
| Q5 | 95.7% (n=1) | 80.6% (n=1) |
| Q9 | 91.0% (n=5) | 77.0% (n=3) |

* at least 100 eggs in each batch.

**Table S3.**
Spo11 mutant cRNA and PCR primer sequences.

| Name | Type | Sequence | Exon targeted |
|---|---|---|---|
| Spo11_exon1_cr5 | crRNA | CAAAGAATAAAAGAGAAACAAGG | 1 |
| Spo11_exon2_cr5 | crRNA | AATTTATAAGAGGCTATTGGAGG | 2 |
| Spo11_exon1_F | Forward Primer | CTCAAACGAAGCTGCGATATTTGCA | 1 |
| Spo11_exon1_R | Reverse Primer | AGCTTCGTGAGAGAGTGCTTTTGAA | 1 |
| Spo11_exon2_F | Forward Primer | GGAGACCTGGATTCGGATTTGGATA | 2 |
| Spo11_exon2_R | Reverse Primer | AGAATGTCTCGGAATGCACAACATG | 2 |

**Data S1. (separate file)**
Tsv file with counts of Mlh1 foci in wildtype nuclei

**Data S2. (separate file)**
Tsv file with raw counts of fertilized and unfertilized wildtype and Spo11 mutant oocytes, as well as number of planula counted from the same experiment

**Data S3. (separate file)**
Zipped folder containing amino acid sequences, alignments, and phylogenetic analyses for CenH3 genes.

**Data S4. (separate file)**
Zipped folder containing amino acid sequences, alignments, and phylogenetic analyses for Mlh1 genes.

**Data S5. (separate file)**
Zipped folder containing amino acid sequences, alignments, and phylogenetic analyses for Piwi genes.

**Data S6. (separate file)**
Zipped folder containing amino acid sequences, alignments, and phylogenetic analyses for Rad51/DMC1 genes.

**Data S7. (separate file)**
Zipped folder containing amino acid sequences, alignments, and phylogenetic analyses for Spo11 genes.

**Data S8. (separate file)**
Zipped folder containing amino acid sequences, alignments, and phylogenetic analyses for Sycp1 genes.

**Data S9. (separate file)**
Zipped folder containing amino acid sequences, alignments, and phylogenetic analyses for Sycp3 genes.
